# Dense encoding of developmental regulatory information may constrain evolvability

**DOI:** 10.1101/2020.04.17.046052

**Authors:** Timothy Fuqua, Jeff Jordan, Maria Elize van Breugel, Aliaksandr Halavatyi, Christian Tischer, Peter Polidoro, Namiko Abe, Albert Tsai, Richard S. Mann, David L. Stern, Justin Crocker

**Affiliations:** European Molecular Biology Laboratory, Heidelberg, Germany; Collaboration for joint PhD degree between EMBL and Heidelberg University, Faculty of Biosciences; Janelia Research Campus, 19700 Helix Dr, Ashburn, VA 20147, USA; Columbia University, Jerome L. Greene Science Center Quad 9A, MC9891 New York, NY 10027

## Abstract

Gene regulatory changes underlie much of phenotypic evolution. However, the evolutionary potential of regulatory evolution is unknown, because most evidence comes from either natural variation or limited experimental perturbations. Surveying an unbiased mutation library for a developmental enhancer in *Drosophila melanogaster* using an automated robotics pipeline, we found that most mutations alter gene expression. Our results suggest that regulatory information is distributed throughout most of a developmental enhancer and that parameters of gene expression—levels, location, and state—are convolved. The widespread pleiotropic effects of most mutations and the codependency of outputs may constrain the evolvability of developmental enhancers. Consistent with these observations, comparisons of diverse drosophilids reveal mainly stasis and apparent biases in the phenotypes influenced by this enhancer. Developmental enhancers may encode a much higher density of regulatory information than has been appreciated previously, which may impose constraints on regulatory evolution.

**Quote:** “Rock, robot rock

Rock, robot rock

Rock, robot rock”

*Daft Punk (2005)*

## Introduction

Developmental enhancers specify the times, levels, and locations of gene expression and are the substrate for much of phenotypic evolution^1,2^. The primary sequences of orthologous enhancers can be highly divergent, even though they often retain conserved functions^3^. This observation has led to the common assumption that the distribution of regulatory information is distributed sparsely within most developmental enhancers, and that function is conserved through the turnover of binding sites that confer similar regulatory outputs^4,5^.

However, enhancer architecture has been studied primarily through approaches biased towards candidate regions identified genetically, biochemically, or through phylogenetic footprinting^6^. Such biased approaches may have limited our understanding of regulatory logic^7^. Our lack of understanding is highlighted by the fact that even the most well-studied developmental enhancers cannot yet be built synthetically^7,8^. Furthermore, while we now have many examples of how individual nucleotide changes have altered enhancer function during evolution^9–15^, we have almost no understanding of what kinds of mutational changes are evolutionarily accessible or how the distribution of transcription factor binding sites may constrain enhancer evolvability.

Mutational scanning experiments provide an unbiased survey of regulatory inputs ^16–20^, can detect weak regulatory interactions critical for robust and precise expression ^21–23^, and can identify mutational effects on phenotypic plasticity^24^. However, it has been difficult to perform mutational scanning of developmental enhancers, where the precise expression patterns across fields of cells must be examined. To facilitate mutational scanning experiments of developmental enhancers, we developed an automated robotics pipeline that facilitates quantitative measurement of expression levels and patterns for multiple embryonic stages and used it to mutationally scan a developmental enhancer.

## Results

### Mutational scanning of *E3N* footprints enhancer activity

To examine the distribution of regulatory information within a developmental enhancer, we selected an enhancer of the *D. melanogaster shavenbaby (svb)* gene*, E3N,* which drives expression in ventral stripes of the embryo and is required to drive cell differentiation of the ventral denticle belts^25^ (Fig. 1a). We focused on this transcriptional enhancer because (1) the function of this enhancer is largely conserved across species, yet the primary sequence has diverged considerably^26^, and (2) it is only 292 bp long, yet integrates patterning information from multiple patterning networks (Fig. 1b)^26,27^ (Fig. 1c).

**Figure 1.**
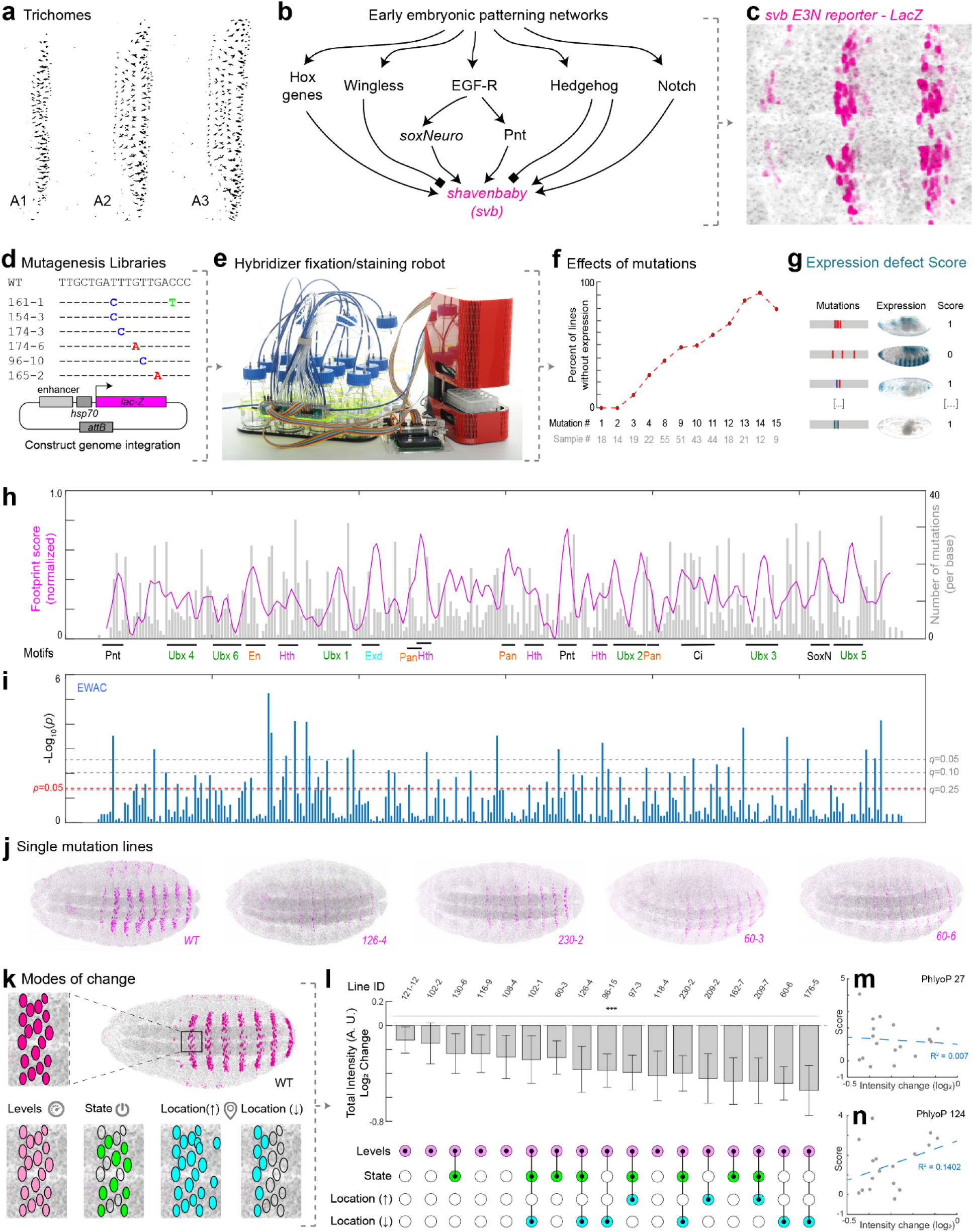
Most nucleotide mutations in *E3N* alter gene expression. (**a**) Laval cuticle prep for *D. melanogaster* for the ventral abdominal segments. (**b**) Schematic of *shavenbaby* regulatory inputs. (**c**) The *shavenbaby WT E3N::lacZ* reporter construct stained with anti-β-Galactosidase. (**d**) Mutagenesis libraries of *E3N* noting mutations and a reporter construct schematic used for integration into the library into the *Drosophila* genome. See also Fig. S1. (**e**) The Hybridizer liquid-handling robot. See also Fig. S2. (**f**) Plot comparing the percent of lines with no expression versus the number of mutations. (**g**) Lines were scored as “1” or “0”, for positive or no visible expression defects, respectively. See Materials and Methods. (**h**) Plot of footprinting scores versus *E3N* sequence. Magenta is the footprinting score (*σ_i_*, see methods). The higher the peak, the higher probability that a mutation will change expression. Gray plots are the mutation coverage for the number of lines screened per base (*M_i_*, see methods). (**i**) EWAC scores represent p-values from a log of odd ratio test for the association of a mutation changing expression. Dashed lines denote p- and q-values ^68^, respectively. See Materials and Methods. (**j**) A subset of the single-mutant *E3N::lacZ* reporter constructs. (**k**) Schematic of possible changes to enhancer outputs. (**l**, top) Single mutation individual nuclear intensity changes compared to WT *E3N* (n = 212 nuclei; N = 8 embryos). In plots, line is mean, upper and lower limits are standard deviation. A two-tailed t-test was applied for each individual comparison. Stars denote p < 0.0001. See methods for sample sizes. AU, arbitrary units of fluorescence intensity. (**l**, bottom) Expression output changes for the single mutations. See also Fig. S3. (**m**) Pearson correlation between mutation effect-sizes and PhyloP scores for 27 species. (**n**) Pearson correlation between mutant effect-sizes and PhyloP scores for 124 species. Regression lines and R-squared values were gathered using The Method of Least Squares.

We generated mutagenized libraries of the *E3N* enhancer with an error-rate of ~2% per mutant, mimicking the divergence in this enhancer between the sibling species *D. melanogaster* and *D. simulans* (Fig. 1d; Fig. S1). We identified mutations in 272 of 292 sites, and the number of mutations per enhancer was approximately Poisson distributed. These enhancers were combined with a heterologous *hsp70* promoter because this enhancer-promoter combination provides a large dynamic range^28^, with the caveat that this design may discount regulatory information that acts only upon the native *svb* promoter. Enhancer variants were integrated as reporter constructs into the *Drosophila* genome at the same genomic location and we isolated 749 lines containing seven mutations on average per enhancer, with a range between one and nineteen (Fig. S1C).

To automate embryo handling and staining, we engineered tools for egg-collections that interface with a chemical handling robot (Fig. 1e; Fig. S2). The “Hybridizer” robot automates several critical steps such as embryo fixation, vitelline membrane removal, and antibody or chemical staining of embryos, allowing classical *Drosophila* fixation and immunohistochemistry protocols on 24 embryonic samples per experiment-providing increased throughput and reproducibility.

To estimate the fraction of sites in this enhancer that confer function, we examined enhancer activity across a subset of 274 lines with variable numbers of mutations by staining embryos carrying enhancers driving a lacZ gene using a Beta-Galactosidase assay. We found that the proportion of lines without any detectable *E3N* expression increased monotonically with the number of mutations in *E3N* (Fig. 1f), suggesting that the *E3N* enhancer contains a high density of regulatory information.

To explore further how this information is distributed, we performed computational “footprinting” to identify important regions within the enhancer. Each mutated base in a line was given a score of 0 if there was no effect on expression, or 1 if the expression was either completely lost or the expression pattern did not resemble that of the wild-type (Fig. 1g). The total score (sum of all lines) for each base was normalized to the total number of mutations per base, smoothed, and plotted (Fig. 1h, see methods). A limitation of this approach is that most lines harbor multiple mutations, and thus most sites were not assayed individually, and the assay is qualitative. Nonetheless, we identified regions that had strong effects on expression, many of which overlapped with previously identified transcription factor binding sites or with consensus motifs for transcription factors.

We next calculated odds ratios across the screened lines to see how the variants were associated with the loss of expression. This approach, which we term <u>E</u>nhancer <u>W</u>ide <u>A</u>ssociation <u>C</u>atalogue (EWAC)^29^, follows a logic similar to a Genome-Wide Association Study^30^. In this case, we used the enhancer-wide set of genetic variants from our library to test whether any individual variant is associated with the loss of enhancer activity more often than expected by chance. We found that many regions were significantly associated with changes in *E3N* regulatory activity when mutated (Fig. 1i), consistent with the density of the footprinting scores. Together, these results suggest that most sites in the *E3N* enhancer are required to generate a wild-type expression pattern.

### Most nucleotide mutations in *E3N* alter gene expression

While the Beta-Galactosidase assay surveyed general trends, it did not allow quantification of subtler differences in enhancer function. We, therefore, developed an imaging pipeline for embryos stained with fluorescent antibodies (Fig. S2). Cleared embryos were imaged using an adaptive feedback confocal microscope pipeline^31,32^. Images were subsequently compiled for analysis as montages or registered using internal fiduciaries (Fig. S3).

To examine the quantitative effects of *E3N* mutations on gene expression, we first screened lines carrying single mutations in *E3N*. A subset of the single mutations is shown in Fig. 1j, demonstrating the substantial effects of most mutations. We next examined possible modes of transcriptional outputs for lines with single mutations: changes in levels, state, and location (Fig. 1k). Previous studies have suggested that each of these modes can be altered independently^33^. In contrast, we found that the majority of *E3N* mutations that caused changes in the levels of expression covaried with changes in transcriptional states and locations (Fig. 1l).

To test whether evolutionary conservation of DNA sequences provides a reliable signal of enhancer site function, we tested for a correlation between the expression changes inferred for single *E3N* sites and the level of conservation for these sites. If evolutionary conservation provided a strong signal of function, then we would expect a positive correlation between expression changes and evolutionary conservation. In contrast, we found no correlation between expression changes and conservation using a *phyloP* estimate of evolutionary conservation among 27 insect species (R^2^ = 0.007) (Fig. 1m), and a weak inverse correlation using a *phyloP* estimate among 124 insect species (R^2^ = 0.140)^34–36^ (Fig. 1n). These results suggest that sequence conservation is a poor predictor of the quantitative roles of individual sites in the *E3N* enhancer (Fig. S3).

Three lines of evidence indicate that regulatory information is encoded densely within the *E3N* enhancer: (1) the number of lines without expression increases as a function of the number of mutations per line; (2) mutational scanning footprinting and EWAC analysis identified many sites required for normal expression; and (3) most single point mutations generated quantitative differences in expression. Based on a random sampling of mutations across the *E3N* enhancer, few sites appear to be non-functional.

### Mutational scanning identifies a Hth binding site associated with a changed evolutionary phenotype

We next sought to validate these results by investigating one site with large effects on expression. We selected eight lines from the library carrying mutations in a Homothorax motif with a high footprint and EWAC score *(Hth2),* and no more than two other mutations elsewhere in the enhancer (Fig. 2a and b). Even though most lines carried different mutations in *Hth2,* most of these lines drove expression in just a single row of cells. We generated targeted knockouts of the *Hth2* and other Hth motifs (Fig. S4a-p). The *Hth2* targeted knockout exhibited low expression levels and expression in fewer cells, similar to the mutant lines in our library (Fig. S4g, h). We validated these results with electromobility gel-shift assays (EMSAs) across the *E3N* enhancer (Fig. S4q-y). The EMSAs demonstrated that Hth and its cofactor Extradenticle (Exd) bind to the *E3N* Hth motifs *in vitro,* while a variant Hth lacking a homeodomain (HthHM/Exd) does not (Fig. S4s). Finally, we found that Ubx binding is enhanced with full-length Hth (Fig. S4t), suggesting possible cooperativity between Hth and Ubx that requires the *Hth2* site (Fig. 2c).

**Figure 2.**
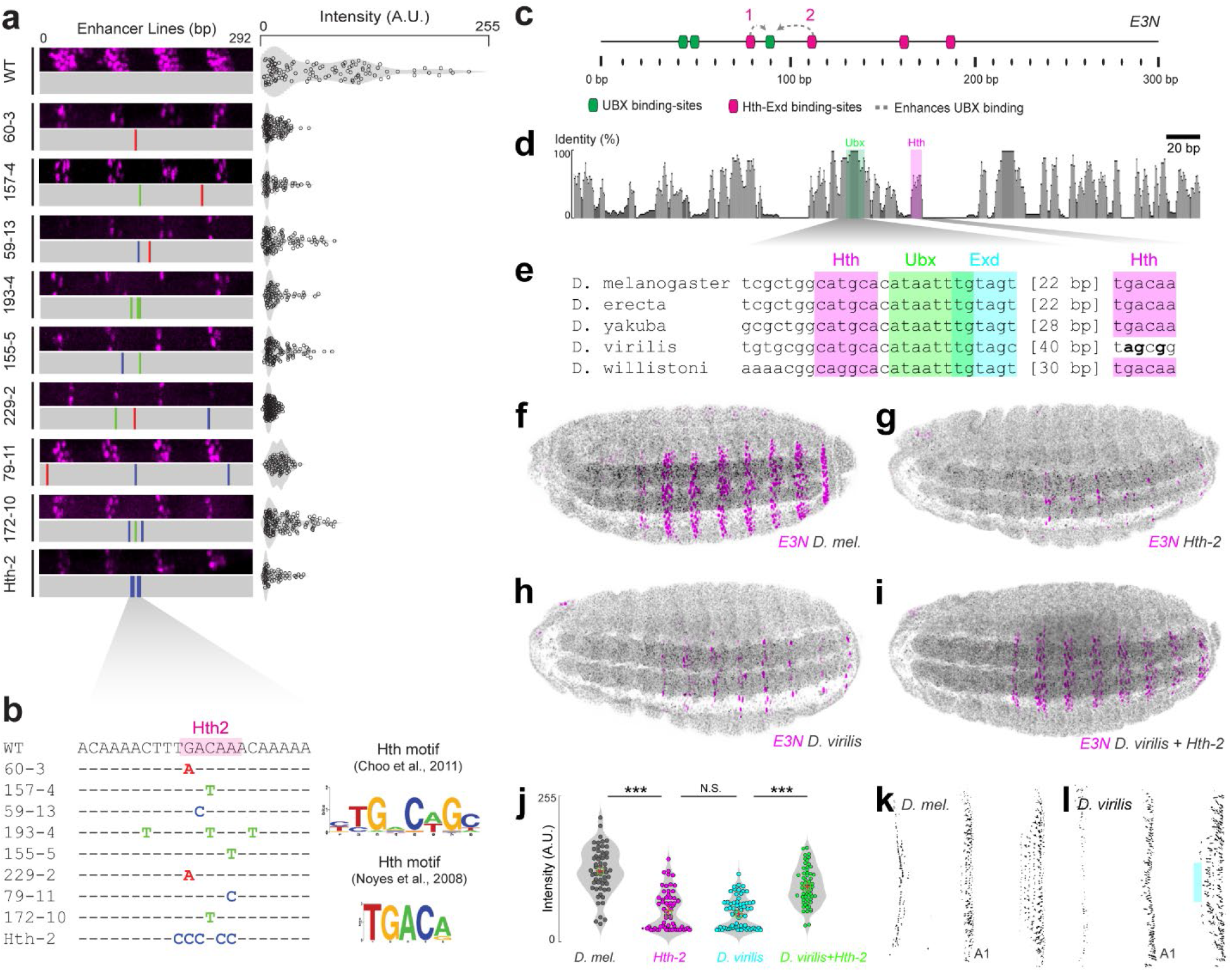
Mutational scanning identifies a Hth binding site associated with a changed evolutionary phenotype. (**a**) Lines tested that contain mutations within a Homothorax binding site (Hth2). Cell-by-cell quantification of the staining intensities in cells with the indicated reporter construct. (**b**) Sequences of lines tested, and sequence logos for characterized Hth binding motifs^69,70^. (**c**) Schematic for the *E3N* enhancer, denoting binding sites and possible protein-to-protein interactions. (**d**) Conservation plot of *E3N* enhancer across 12 *Drosophila* species, highlighting conserved Ubx and Hth binding sites. (**f**-**i**) *E3N::lacZ* reporter constructs in a *Drosophila melanogaster* background stained with anti-β-Galactosidase. (f) *D. mel* WT *E3N::lacZ* reporter construct. (g) *D. mel E3N::lacZ* reporter construct with a mutated Hth2 site (TGACAA ➔ CCCCCC). (h) *D. virilis E3N::lacZ* reporter construct. (i) *D. virilis E3N::lacZ* reporter construct with rescued Hth2 site (TAGCGA ➔ TGACAA). (j) Cell-by-cell quantification of the staining intensities in cells with the indicated reporter construct. Mean and median are shown as red crosses and green squares, respectively. In box plots, center line is mean, upper and lower limits are standard deviation and whiskers show 95% confidence intervals. Two-tailed t-test was applied for each individual comparison, stars denote p < 0.01 (n = 50, 10 embryos). (k and l) Cuticle preps for *D. mel* (k), and *D. virilis* (l) showing ventral segments. Teal overlay highlights band of trichomes not present in *D. virilis.*

Despite the importance of the *Hth2* binding site in the *D. melanogaster E3N* enhancer, this site is not strongly conserved between different species of *Drosophila* and is not present in *Drosophila virilis* (Fig. 2d, e). In contrast, other sites, such as Hth1 and Exd, are conserved (Fig. 2e; Fig. S5). A stable transgenic *vir-E3N* reporter in *D. melanogaster* drove an almost identical expression pattern as the *mel-E3N Hth2* deletion, and both enhancers drove lower expression than the wild type *mel-E3N* enhancer (Fig. 2f-h).

To test if the loss of the *Hth2* site contributes to the weaker expression of the *vir-E3N* enhancer, we “resurrected” the *Hth2* motif in the *vir-E3N* enhancer (Fig. 2i). The addition of this binding site increased the expression of the *vir-E3N* to nearly the level of the *mel-E3N* enhancer (Fig. 2j). Finally, we found that *D. virilis* larvae exhibit fewer ventral trichomes in the most anterior ventral band of denticles (Fig. 2k, l), the domain in which *mel-E3N* is active, suggesting that the loss of *Hth2* in the *D. virilis E3N* enhancer may have contributed to the loss of ventral trichomes in *D. virilis.*

### Mutations in Ubx binding sites often simultaneously altered levels, timing, and locations of *E3N* expression

The *E3N* enhancer appears to contain regulatory information in most nucleotide positions. One consequence of this high density of regulatory information is that it may limit the possible expression patterns that can be generated through stepwise mutation and may bias change in some directions^37–39^. We examined this question first by analyzing the manifold consequences of mutations in Ubx binding sites. In *E3N*, Ubx acts as a transcriptional activator and we previously identified multiple low-affinity Ubx binding sites in this enhancer^26,40^.

An important feature of our pipeline is that it captures a range of *Drosophila* embryo stages. We found that the higher-affinity Ubx motif drove earlier expression, activating the enhancer prematurely in stage 14 embryos (Fig. 3a-d). Additionally, the intensities in both anterior, early stripe, and naked (inter-stripe) regions in the Ubx High-Affinity line were significantly higher at stages 14 and 15 than for the wild type *E3N* enhancer (Fig. 3e).

**Figure 3.**
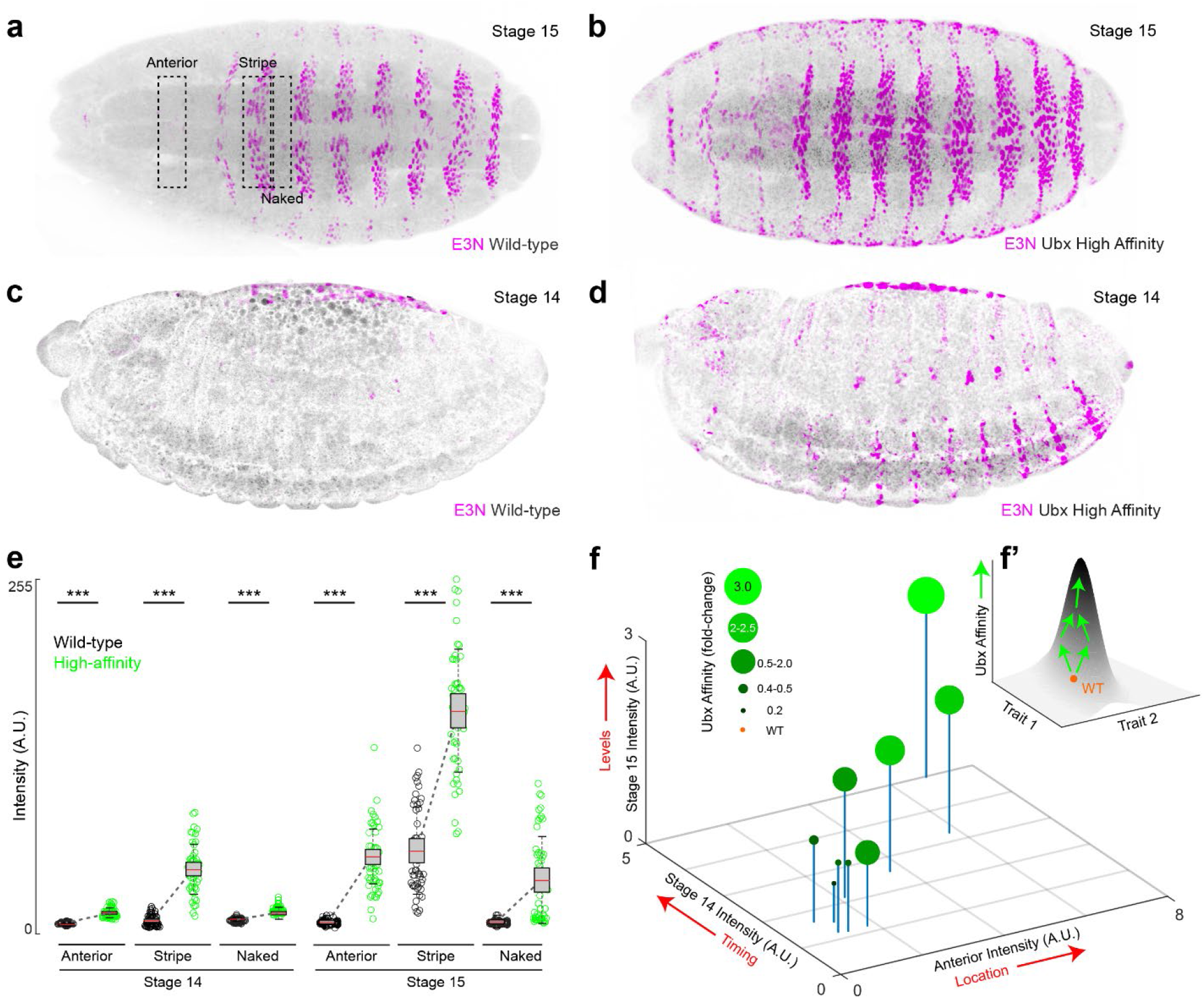
Ubx mutations often simultaneously change levels, timing, and locations of *E3N* expression. (**a-d**) *E3N::lacZ* reporter constructs at stages 14 and 15 stained with anti-β-Galactosidase. (**a** and **c**) WT *E3N::lacZ* at stages 15 and 14. (**b** and **d**) *E3N::lacZ* construct with increased Ubx affinity at stages 15 and 14. (**e**) Measurements of stripe (intra) and naked (inter-stripe) expression and ectopic anterior expression during stages 14 and 15. Cell-by-cell quantification of the staining intensities in cells with the indicated reporter construct. In box plots, center line is mean, upper and lower limits are standard deviation and whiskers show 95% confidence intervals. Two-tailed t-test was applied for each individual comparison, stars denote p < 0.01 (n = 50, 10 embryos). (**f**) Plot of measurements between different mutant lines comparing total Ubx affinity (shades of green and circle size), anterior intensity, stage 14 intensity, and stage 15 intensity (**f**). (**f’**) Model for Ubx affinity and linked traits.

We next wanted to determine if these trends extended across a wider range of Ubx affinities. Using NRLB^40^ to calculate the total Ubx affinity across the mutation library, we selected a subset of lines with varying total Ubx affinities, with the number of mutations outside the Ubx binding sites ranging from 0 to 3. We analyzed the anterior, stripe, and naked (inter-stripe) region intensities for each of the lines and plotted the results (Fig. 3f). We found that these phenotypes were correlated and expression levels within all domains of expression increased with increased Ubx affinities (Fig. 3f, f’). Strikingly, this increase appears to be coupled to (1) precocious expression, appearing a stage earlier than usual, and (2) ectopic expression in naked regions and anterior domains. These results could be explained by the enhancer having a higher sensitivity to Ubx levels, resulting in precocious expression, which is also linked to ectopic expression resulting from the binding of other homeodomain factors to higher-affinity sequences^25,26^.

To explore whether the pleiotropic effects observed for Ubx—which acts as an activator on this enhancer—generalized to other sites, we next examined binding motifs for a transcriptional repressor. The Wingless/Wnt signaling pathway represses *svb* expression in naked regions between denticle bands on the ventral larval cuticle^27,41^ (Fig. 4c). Disruption of *wingless (wg)* function leads to Svb overexpression, resulting in a lawn of trichomes on the larval cuticle (Fig. 4a, b), demonstrating that it is *possible* to drive ectopic trichomes between rows of denticles.

**Figure 4.**
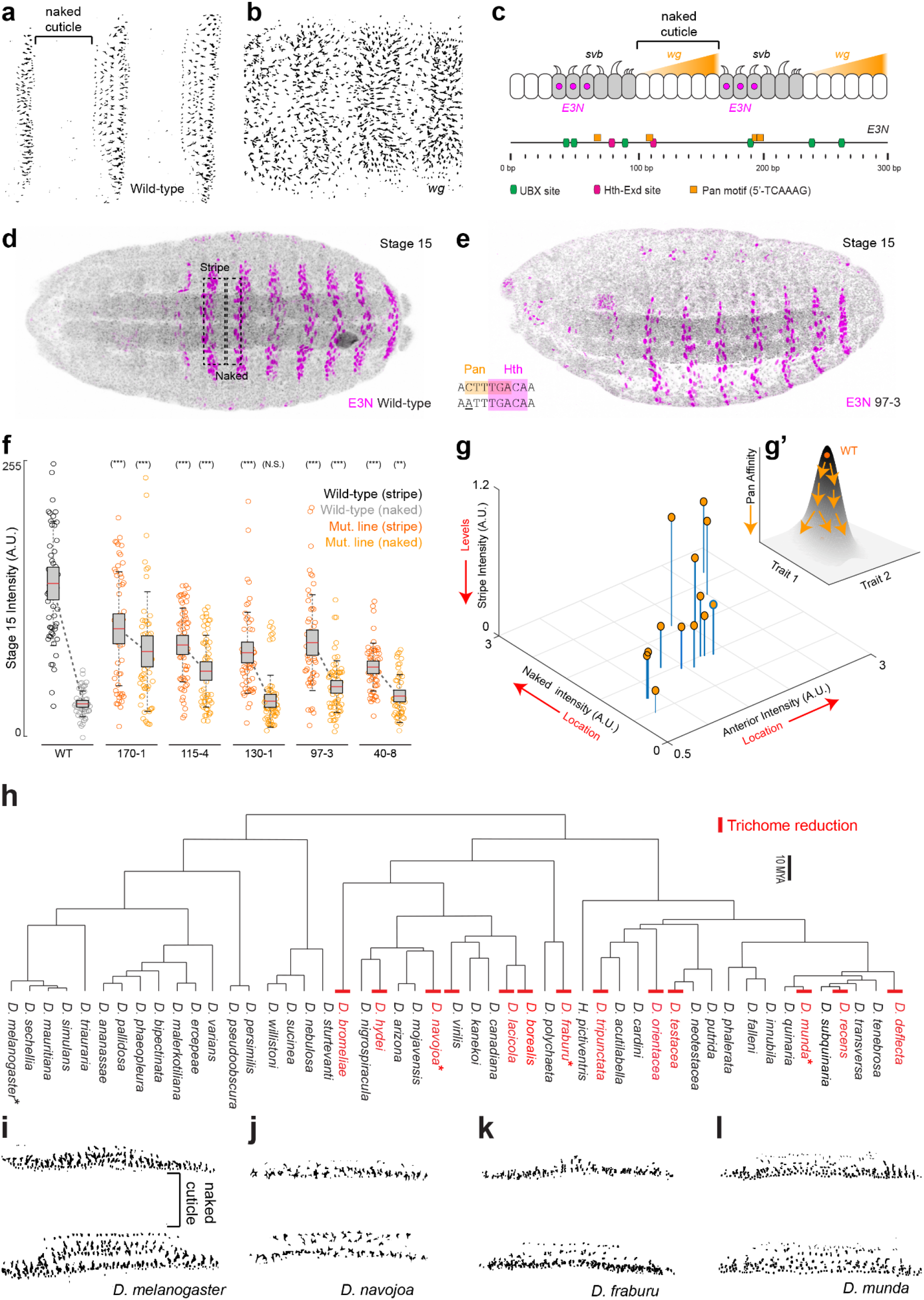
*E3N* enhancer architecture may constrain evolutionary paths. (**a**) WT (*w^1118^*) cuticle prep. (**b**) *wg* deficient background *(wg^CX4^)* cuticle prep. (**c**) Top: model of *Wg* expression in naked cuticle regions that represses interstripe activity through Pan (Tcf) activity. Bottom: schematic of the *E3N* enhancer denoting motifs for activators (Ubx and Hth) and repressor motifs (Pan). (**d** and **e**) *E3N::lacZ* reporter constructs stained with anti-β-Galactosidase for the wild-type enhancer (**e**) and the mutation line *97-3::lacZ* (**e**) See also Fig. 2G. (**f-g’**) Boxplots comparing expression in the naked and stripe regions across a subset of the Pan mutation lines (**f**). Cell-by-cell quantification of the staining intensities in cells with the indicated reporter construct. In box plots, center line is mean, upper and lower limits are standard deviation and whiskers show 95% confidence intervals. Two-tailed t-test was applied for each individual comparison, stars denote p < 0.01 (n = 50, 10 embryos). (**g**) Lines plotted comparing intensities in inter- and –intrastripe regions with anterior expression, and a schematic (**g’**) for the relationship of Pan affinity and traits lowering total expression when increasing ectopic expression. (**h**) A *Drosophila e*volutionary tree noting gains (no examples) and losses (red) of trichomes in the naked region. Asterisks denote cuticle preps highlighted in panels (**i-l**). Cuticles preps from *D. melanogaster* (**i**), *D. navojoa* (**j**), *D. fraburu* (**k**), and *D. munda* (**l**), see also Fig. S6.

The Wingless responsive transcription factor Pangolin (Pan) binds to known DNA motifs^42^. We selected lines from our library carrying mutations in Pan motifs, including a line with a single mutation overlapping with the previously characterized *Hth2* site and a Pan motif (Fig. 4d, e). Across the 13 lines carrying mutations in Pan motifs, we found both lower levels of expression in the denticle “stripe” domain and ectopic expression in the naked region (Fig. 4f). Increased ectopic expression was negatively correlated with overall expression levels (Fig. 4g, g’). Together, these observations of the effects of changes in activator and repressor binding sites demonstrate that different aspects of enhancer expression may be correlated because of the pleiotropic effects of single binding sites.

### *E3N* enhancer architecture may constrain evolutionary paths

The widespread pleiotropic effects of mutations across the *E3N* enhancer suggest that the distribution of regulatory information may constrain evolutionary paths for this enhancer^39^. We, therefore, examined larval cuticles of 60 *Drosophila* species (Fig. 4h, Fig. S6) to see whether the ventral larval trichomes, which result in part from the activity of *E3N*, show patterns of constraint across the genus *Drosophila.* While genetic perturbations can lead to the development of trichomes in the canonical inter-stripe domain (see Fig. 4b), we did not observe trichomes in this anatomical domain in any of the species (Fig. 4h-l). Furthermore, we detected at least twelve evolutionary losses of trichomes within ventral denticle bands. Together, these results are consistent with the hypothesis that expression driven by the *svb E3N* enhancer is constrained across *Drosophila,* and that changes in *E3N* may more often lead to loss of trichomes than to gain in new anatomical domains.

## Discussion

We have illustrated an unbiased method to explore regulatory regions in developmental systems. One striking result of our experiments is that most mutations in the *E3N* enhancer led to changes in transcriptional outputs, suggesting that regulatory information is distributed densely within this enhancer. The high density of regulatory information in this enhancer, combined with the pleiotropic effects of mutations in many sites, may constrain enhancer evolution. These observations are surprising because the DNA sequence of *E3N*, like most developmental enhancers, evolves rapidly although the function is largely conserved^1^.

Our results are similar to those found for a minimal 52-bp rhodopsin promoter, where 86% of single nucleotide substitutions significantly altered expression^35^, suggesting that dense regulatory logic may be a general feature of regulatory regions. This may be why it has been so hard to predict and to build synthetic enhancers^7^.

One explanation for both the observed fragility of *E3N* and its rapid evolution is that *E3N* has evolved along constrained mutational paths^39, 43^. However, there are several other possible explanations. First, removing an enhancer from its native location and placing it in a novel chromatin environment may facilitate binding of transcription factors to sites that are not used in their native context^44^. Second, changes in binding sites may normally be buffered by regulatory information adjacent to the enhancer in its native context^45,46^. Finally, additional buffering can result from regulatory information encoded by partially redundant enhancers^47–50^. Thus, isolated enhancers may be fragile, but in their natural context, additional regulatory information could hide the functional consequences of changes in individual enhancers, ultimately allowing relatively rapid evolution of enhancers while retaining the overall function of the gene.

We observed that all mutations that generated ectopic expression in naked regions exhibited pleiotropic ectopic expression in other domains. Thus, it appears that there are no mutations in the lines we tested that enable *E3N* to escape this pleiotropic constraint. Consistent with this observation, we did not observe any *Drosophila* species that exhibited trichomes in these domains. Although it is also possible that natural selection has not favored the presence of trichomes in these domains, this potential regulatory constraint may help explain the absence of trichomes in inter-stripe regions in the genus *Drosophila.* Other studies have shown that mutational changes in other *svb* enhancers, which result in altered dorsal trichome patterns^51^, tend to act pleiotropically across most segments of the larval body, suggesting that multiple *svb* enhancers are mutationally constrained. Similarly, another *svb* enhancer (*E6*) encodes extensive redundancy of activator binding sites, which may have biased evolution toward the gain of a novel binding site for a strong repressor to allow escape from the constraint of redundancy^9^.

There are additional sources of enhancer constraint. For example, the *E3N* enhancer encodes low-affinity binding sites that confer specificity for a subset of Hox proteins^26^. Low-affinity binding sites facilitate the production of precise temporal and spatial patterns of expression^21,52^. This requirement for low-affinity binding sites, however, means that the entire enhancer may require many transcription factor binding sites, and a specific arrangements of binding sites^53,54^, to preserve kinetic efficiency^21,26,40^. In the case of the *E3N* enhancer, cooperativity between Hth and Ubx may enable the enhancer to drive precise patterns of expression but make it fragile to mutations. Finally, spatial precision can be encoded through binding site competition between transcriptional activators and repressors^33^, consistent with the overlapping Pan/Hth motifs. It may be hard to separate the regulatory inputs of either factor independently through binding site turnover. This codependency may also constrain evolvability because modification of overlapping sites leads to pleiotropic effects. Thus, developmental constraints may operate on enhancers through multiple mechanisms, explaining the deep conservation of some elements^55–57^.

In the future, we hope that our pipelines will serve as a platform for future research by allowing a broader community of users to build, execute, and share similar experiments. This approach to studying enhancer function is a powerful way to explore modes of regulatory evolution that consider both “the possible and the actual” operating on regulatory regions^58^, and may, therefore, allow us to predict changes in an evolutionary context.

## Supporting information

Library Sequences

## Acknowledgments

Albert Tsai is supported by the German Research Foundation (Deutsche Forschungsgemeinschaft, TS 428/1-1) and EMBL. We thank J. Zaugg and J. Wirbel for statistical advice; CJ. Standley, X. Li, RM. Galupa, GA. Canales, and MRP. Alves for helpful discussions with the manuscript; C. Rastogi and H. Bussemaker for the NRLB algorithms; K. Richter and J. Sager for technical support; T. Laverty with *Drosophila* assistance; and A. Milberger with CAD design. We thank Joshua Payne for insightful discussion.

## Author contributions

T.F., M.B. and J.C. performed the measurements, R.M., D.S. and J.C. were involved in planning and supervised the work, T.F. and J.C. processed the experimental data, performed the analysis, drafted the manuscript and designed the figures with input from all the authors. J.J., and P.P. built the robotics with input from J.C. A.H., and C.T. write the automated microscopy code and analysis tools. N.A carried out the biochemistry experiments. T.F., A.T., and J.C. analyzed the data.

## Data and material availability statement

The original images (cuticle preparations and embryo images, organized into zip files) will be available for download and are indexed at: https://www.embl.de/download/crocker/svb_E3N_screen/index.html. All fly lines will be available upon reasonable request.

## Materials and Methods

### Fly strains and constructs

All reporter constructs and the *E3N* mutant library were synthesized and cloned (GenScript) into the *placZattB* reporter construct. All mutant genetic sequences are available in Table S1.

### Embryo fixation and robotics

*Drosophila* were loaded into egg collection chambers (Fig. S2). Embryos were collected overnight and washed in a saline solution (0.1 M NaCl and 0.04% Triton X-100), dechorionated in 50% bleach for 90 seconds, and rinsed with water. For manual fixation, embryos were transferred to scintillation vials containing fixative solution (700 uL 16% PFA, 1.7 mL PBS/EGTA, 3.0 mL 100% heptane) and fixed for 25 minutes, shaking at 250 rpm. The lower phase was separated, and embryos were shocked isotonically using 100% methanol and rapid vortexing for 30 seconds. The interphase and upper phase were removed and the embryos were washed in fresh methanol. Fixation and antibody staining was tested on a series of control wells and showed no significant difference between sample fluorescence (Fig. S2)

For automated fixation, embryos were transferred to fixation plates (Fig. S2) and loaded into the robot. Scripts for the Hybridizer, 3D CAD files, and the fixation protocol (reagents same as manual fixation) can be found on Github: (https://github.com/ianelia-pypi/hybridizer_python/tree/digital)

### Antibody staining and cuticle preps

Fixed embryos were stained using standard procedures with a chicken anti-βGal (1:500, abcam ab9361) and mouse anti-ELAV supernatant (1:25, DSHB), and conjugated respectively with AlexaFluor 488 and 633 secondary antibodies (1:500, Invitrogen). *Drosophila* cuticles were prepared using standard protocols^59^. The anti-ELAV antibody was obtained from the Developmental Studies Hybridoma Bank, created by the NICHD of the NIH and maintained at The University of Iowa, Department of Biology, Iowa City, IA 52242.

### Protein Purification and EMSAs

Ubx (isoform IVa), HthHM-Exd, and HthFL-Exd constructs, protein purification, and EMSA conditions were described previously ^26^.

### Embryo mounting

Fixed and antibody-stained embryos were either mounted on Prolong Gold (Thermo Fischer Scientific) or mounted in BABB (benzyl alcohol / benzyl benzoate). For BABB mounting, embryos were serially washed into 100% Ethanol, 2× in BABB, and incubated overnight.

A Grace Silicone 2×4 well Isolator 2×4 (JTR8R-A2-0.5, Gracio Bio-Labs, USA) was cut in half and applied with another Isolator to a 75 x 50 mm microscope slide (Corning, USA). Using a micropipette, 100 uL of the embryos in BABB solution were transferred to each well. The embryos were allowed to sink, and wells were connected with a thin layer of BABB and covered with a coverslip. The coverslip was sealed with 3 coats of clear nail polish.

### Microscopy and data analysis

Cuticle preps were imaged on a phase-contrast microscope (Zeiss, Germany). Confocal images were acquired on a Zeiss LSM 880 confocal microscope (Zeiss, Germany) either manually or using developed adaptive feedback microscopy pipeline (https://git.embl.de/grp-almf/feedback-fly-embryo-crocker). The pipeline processes acquired images during the experiment and guides the microscope via the MyPic macro^60^ to automatically acquire high-zoom images in the identified positions. Low-resolution overview 3D tile scans were acquired using a 5x/0.16NA air objective and used to detect the lateral positions of embryos. For each selected position, low-resolution 3D stacks were acquired with a 20x/0.8NA air objective lens. The low-resolution stacks were automatically analyzed to obtain the embryo’s bounding box. Within the bounding box a multichannel 3D stack was acquired with the same 20x/0.8NA air objective, however, now at high resolution for quantification. Imaging parameters such as lasers, emission filters, step-sizes, etc. can be found in the .lsm files in the provided link above.

Using Fiji (https://imagej.net/Fiji) ^61^, images for each embryo were max-projected and compiled into montages using the *MontageMaker* plugin (Figure S3a). Embryos could also be registered. (Fig. S3b-d) Embryos were rotated in 3D space using a fiduciary stain (ELAV, DSHB). Maximum projections of the ventral half of the embryo were calculated, and multiple images elastically transformed to one another ^62^, creating composite expression patterns.

Phenotypes were analyzed by a sliding window method (Figure S3e), where a box was drawn between the T1 and T2 segments, centered over T2, and the average intensity for the T2 stripe was taken as the ROI was dragged down the embryo. Additionally, a state-measuring method was employed (Figure S3f), where a nucleus-sized circle ROI measured nuclear intensities down a single column of cells. Finally, entire expression profiles in Figure 3 were generated by drawing a single box around the expression pattern and generated using *ProfilePlot* (Figure S3g). Box plots were generated in Matlab using the *notBoxPlot* plugin ^63^, and violin plots for state-measurements were generated in Matlab using *distributionPlot.m* ^64^.

Figure 1 individual nuclei were identified using the automated threshold algorithm on FIJI and a watershed to split large ROIs. Average intensities for each nucleus were measured and plotted. The number of nuclei and embryos from left to right (n = 72, 278, 98, 142, 169, 168, 247, 169, 136, 107, 325, 177, 211, 241, 221, 272, 256 nuclei; N = 4, 10, 7, 7, 8, 10, 10, 10, 8, 5, 10, 10, 8, 8, 8, 8 embryos).

### X-gal Staining and Imaging

Embryos were washed and dechorionated as described in the Embryo Fixation subsection, and fixed in a 1:1 solution of Fixative (2% formaldehyde + 0.2% glutaraldehyde + PBS) and 100% Heptane for 20 minutes, shaking at 200 rpm. Fixative solution and heptane were removed, and embryos were patted dry on paper towels. Embryos were washed 3x in PBT for 10 minutes, shaking at 150 rpm. Embryos were mixed with X-gal staining solution (5-bromo-4-chloro-3-indolyl-β-D-galactosidase [20 mg/mL DMF], 400 mM potassium ferricyanide, 400 mM potassium ferrocyanide, 200 mM magnesium chloride, H2O), incubating at 37°C for 1 hour. Staining was stopped with 3x PBT washes. Embryos were imaged on a Leica DFC420C Digital Camera using a Leica MZ16F Stereomicroscope.

### Calculating Footprinting Scores

Enhancer sequences were aligned to each other using the *pairwise2* alignment function in Biopython ^65^. For this assay, deletions in the sequences were treated as mismatches, and we removed inserted bases for the analysis, making all sequences 292 bps. Alignments were changed to binary values, where 0 = match, and 1 = mismatch.

For each base (*i* = 1…292) in the *E3N* enhancer and each mutagenized line (*j* = 1…274), a score *α_i,j_* = 0 was assigned if the base in the line is not mutated. For a mutated base, *α_i,j_* = 1. The Mutation Coverage *Mi* (see Fig. 1h) is the sum across all 274 lines:

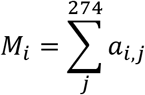

For each base (*i* = 1…292) in the *E3N* enhancer and each mutagenized line (*j* = 1…274), a score *s_i,j_* = 0 is assigned if the base in the line is not mutated or is mutated but did not alter E3N expression pattern. For a mutated base that affected *E3N* expression pattern, *s_i,j_* = 1. The total score *S_i_* at each base *i* is the sum across all lines:

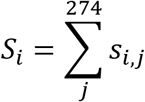

If a specific base is mutated less than 5 times across all lines, the score is discarded (*S_i_* ≡ NaN in the data) and no longer included. A normalized footprint score *σ_i_* is calculated by:

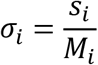

The plot in Fig. 1J is computed in Matlab using the Matlab *smoothdata* function with a Gaussian-weighted moving average window of 5 bases. Excluded bases (*S_i_* = NaN) are ignored by this script as described in its documentation. Footprinting data are available in Table S3.

### Calculating EWAC Scores

For each base (*i* = 1.292) in the *E3N* enhancer, the total score *A_i_* at each base *i* is the total coverage *M_i_* subtracted from *S_i_* (see Calculated Footprinting Scores):

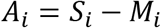

If a value was not available for *S_i_* (*S_i_* = NaN) *A_i_* was set to = 0.5.

The total score *C_i_* at each base (*i* = 1…292) is the score *A_i_* subtracted from the total number of lines without any expression (Q = 129), subtracted from the total number of lines (J = 274):

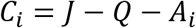

The total score *D_i_* at each base (*i* = 1…292) is the score *S_i_* (see Calculated Footprinting Score) subtracted from the total number of lines without any expression (*Q* = 129):

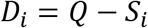

For each each base (*i* = 1…292) a 2×2 contingency table was generated for the values previously calculated:

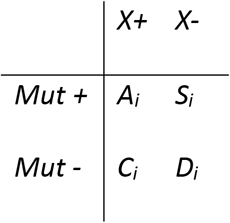

P-values were calculated using *chi2_contingency* from *SciPy*^66^ and compiled using *Pandas*^67^ (see Fig. S1). EWAC scores in Fig. 1 are the –log_10_ of the p-values. Q-values were calculated according Storey et al^68^. EWAC P-values are available in Table S2.

---

**Figure S1.**
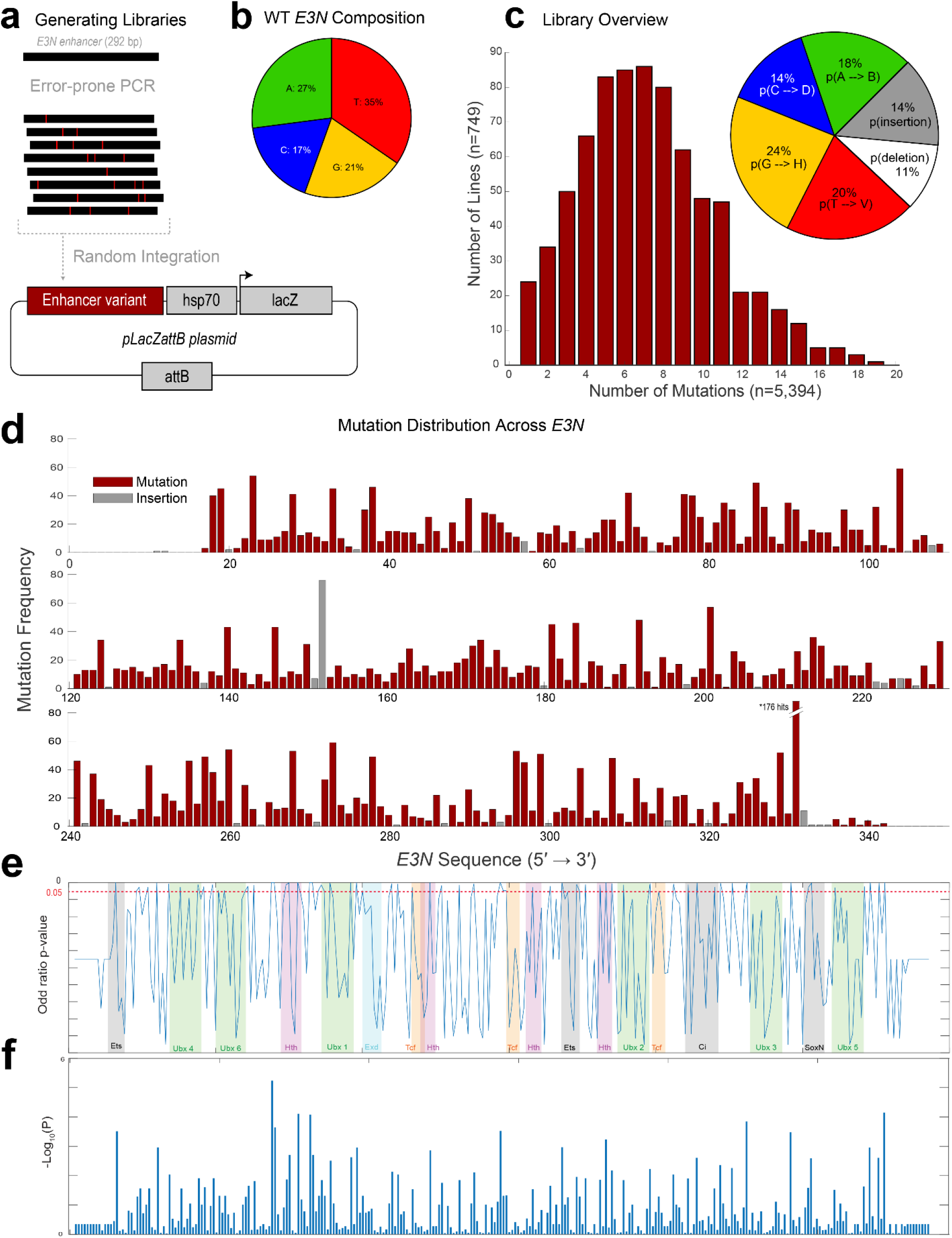
Distribution of mutations in the *E3N* enhancer library. (**a**) Mutant enhancer variants of *E3N* were created via degenerate PCR and integrated into the *placZattB* plasmid, which contains a minimalized core *hsp70* promoter and the *lacZ* reporter gene. Plasmids were integrated into the *Drosophila* genome using at *attP2.* (**b**) Pie chart depicting base-pair composition of the WT *E3N* enhancer. (**c**) (Left) Histogram for all 749 mutants is normal with an average of 7 mutations per mutant. (Right) pie chart shows probability of mutation normalized to ATCG composition from (see b). (**d**) Manhattan plot shows the summation of all mutations within the *E3N* library. (**e**) Calculated log of odds ratios p-values (y-axis) plotted over transcription factor binding motifs (colored and shaded regions) across the *E3N* genomic sequence (x-axis). (**f**) –Log_10_(P) scores plotted as a bar graph.

**Figure S2.**
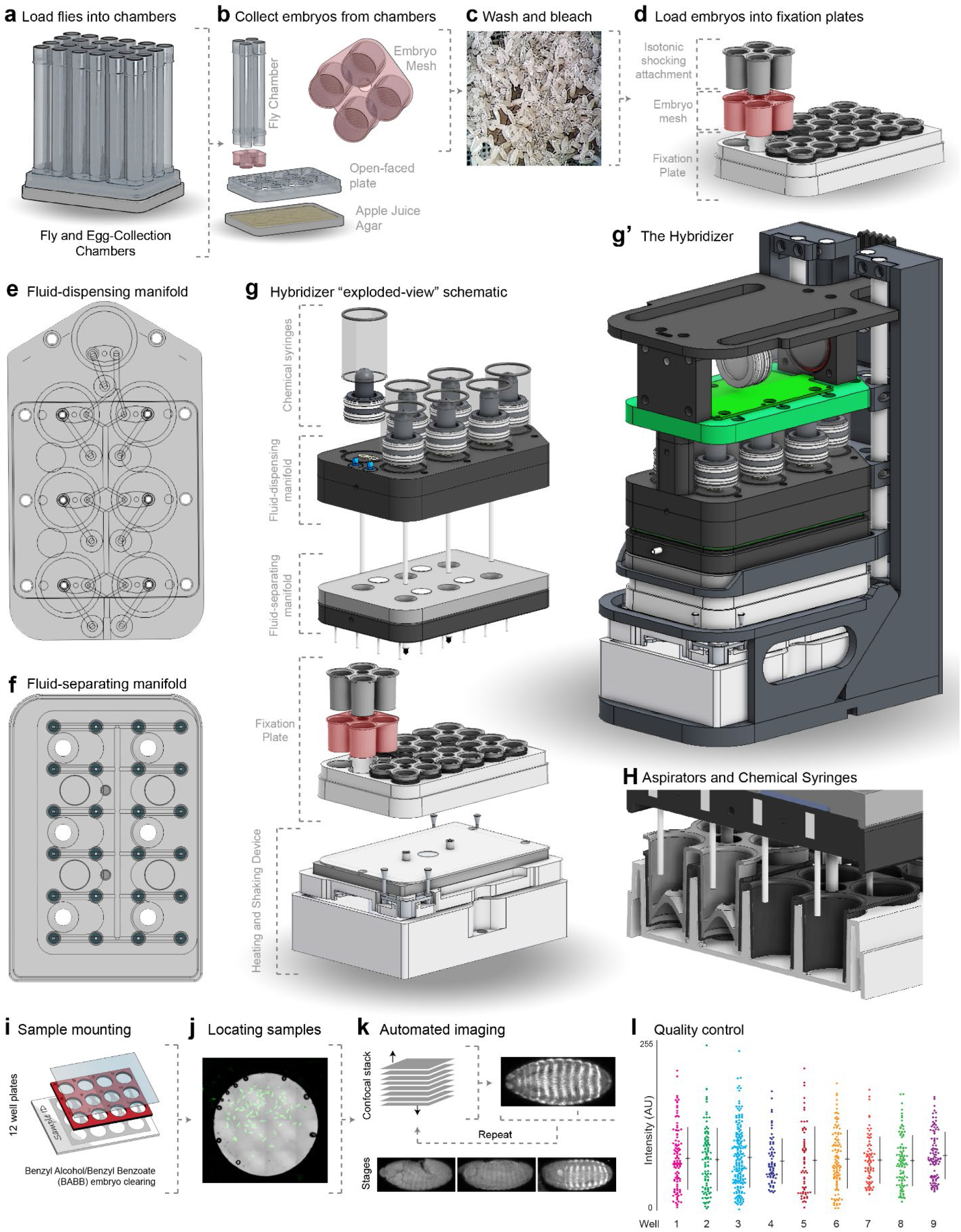
An automated platform for fixing, staining, and imaging *Drosophila* embryos. **(a-d)** Collecting *Drosophila* embryos. (**a**) Custom fly chambers were made, holding up to 24 different strains. (**b**) An explosion-view of the fly chambers. Embryo meshes (red) can attach and detach from the fly chambers and are suspended above an apple juice-agar plate with an open-faced plate. (**c**) Embryos are collected onto the embryo meshes and washed with saline solution and bleached. (**d**) Embryos are loaded into a fixation plate. (**e-h**) Components of the Hybridizer robot. (**e**) The fluid-dispensing manifold. Seven chemical chambers are set on the fluid-dispensing manifold to prime the tubing or dispense chemicals into the fixation plate. (**f**) The fluid-separating manifold uses 24 small syringes to aspirate fluid from the isotonic shocking attachments. (**g** and **g**’) Different components of the Hybridizer. (**h**) Cross-section of the fixation plate and aspirators and syringes. 24 small syringes aspirate fluid from the top of each well within the isotonic shocking attachment and 6 main syringes dispense fluid into the wells and remove fluid from the bottom of the wells. (**i-k**) The adaptive feedback imaging pipeline. (i) Samples are mounted on multi-well slides. (**j**) An overview tile-scan of each well is taken and x,y coordinates for embryos (green) are identified either manually or computationally. (k) For each coordinate, a fast, low-resolution confocal stack is automatically acquired. An algorithm determines the embryo’s z position and rotation, yielding a bounding box within which a high-resolution, 3D stack of the entire embryo is acquired. See also Methods. (**l**) Control *E3N* WT embryos were fixed and stained on the Hybridizer. Nine wells were selected and their fluorescence intensities measured. Cell-by-cell quantification of the staining intensities in cells with the indicated reporter construct. In plots, center line is mean, upper and lower limits are standard deviation. AU, arbitrary units of fluorescence intensity.

**Figure S3.**
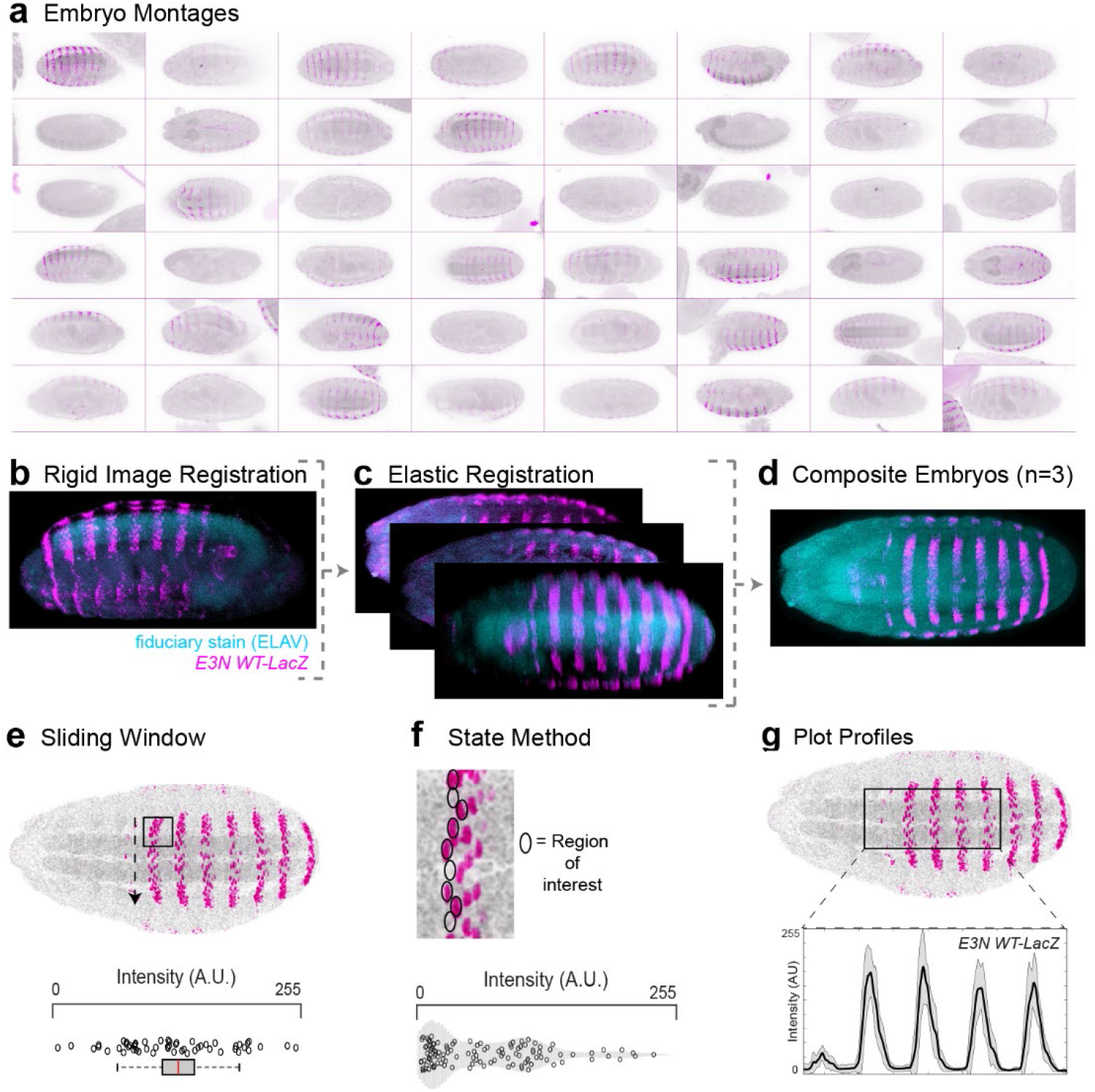
Methods of image and data analysis. (**a**) Images acquired from automated imaging are compiled into a large montage image. (**b-d**) Registering multiple images using fiduciary stains. An embryo acquired during automated imaging (**b**) can be automatically rotated in 3D space using ELAV (teal) as a fiduciary stain. Once properly rotated, maximum projections of the ventral half can be computed (**c**). Finally, the 2-D projections can be elastically registered – or deformed – to align multiple samples (**d**). (**e-g**) Methods of measuring expression patterns. (**e**) Sliding window analysis. A box is drawn between A2 and A3 and centered within A2. Multiple measurements are taken, sliding the box across the stripe. Each point on the boxplot represents one measurement within the box. Cell-by-cell quantification of the staining intensities in cells with the indicated reporter construct. In box plots, center line is mean, upper and lower limits are standard deviation and whiskers show 95% confidence intervals. (**f**) State method analysis. A row of cell-sized ROIs are dragged down across the A2 stripe. Each point on the boxplot represents a single nucleus. (**g**) Plot profile analysis. A box is drawn from the A1 segment to A5 and the mean intensity is taken for each column of pixels and plotted (n = 10). Shaded areas indicate ±1 SD.

**Figure S4.**
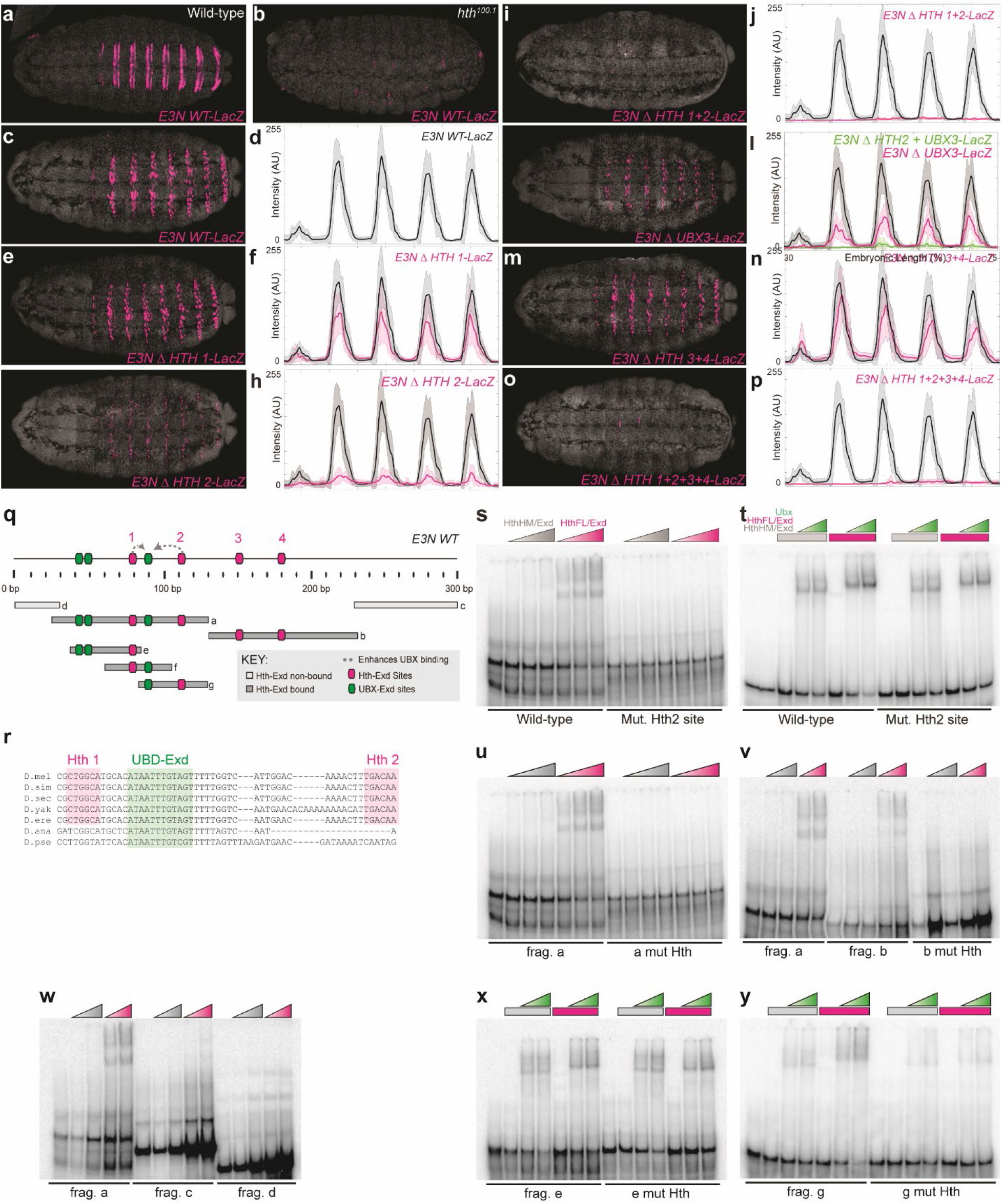
Testing additional Hth-Exd motifs in *E3N*. (**a** and **b**) *E3N::lacZ* reporter constructs in a WT *w^1118^* background (**a**) and *hth* homeodomain-less (HthHM) *hth^100.1^* background stained with anti-β-Galactosidase (**b**). (**c-p**) *E3N::lacZ* reporter constructs stained with anti-β-Galactosidase adjacent to their respective expression plot profiles. Constructs contain mutations to Hth1 (CTGGCA ➔ CCCCCC), Hth2 (TGACAA ➔ CCCCCC), Hth3 (TTGTCG ➔ CCCCCC), and Hth4 (TGAGAG ➔ CCCCCC). (**c** and **d**) *E3N* WT. (**e** and **f**) *E3N* with Hth1 site changed. (**g** and **h**) *E3N* with Hth2 site changed. (**i** and **j**) *E3N* with Hth1 and Hth2 sites changed. (**k** and **l**) *E3N* with Ubx3 site changed (CATAATTTGT ➔ CAGGGTTTGT). (**m** and **n**) *E3N* with Hth3 and Hth4 sites changed. (**o** and **p**) *E3N* with Hth1, Hth2, Hth3, and Hth4 sites changed. In all plots, the black and magenta lines denote the average expression driven by the wild-type and modified enhancers, respectively (n = 10 for each genotype). Shaded areas indicate ±1 SD. AU, arbitrary units of fluorescence intensity. (**q**, top) Schematic for the *E3N* enhancer, denoting binding sites and possible protein-to-protein interactions. (**q**, bottom) Schematic for different *E3N* fragments tested. (**r**) Multiple-species alignment of Hth1, 2 and the UBX-Exd site. (**s-y**) Electromobility shift assays for different fragments of *E3N* denoted in (**m**). HthHM/Exd is the homeodomain-less (HthHM) isoform of Hth incubated with Exd. HthFL/Exd is the Hth isoform with a homeodomain, incubated with Exd. Fragments tested with the WT Hth binding site and a mutated form. (**s**) EMSA for *fragment-f* with Hth2 mutated (**t**) additionally with increasing Ubx concentrations. (**u)** EMSA for *fragment-a* with Hth1 and 2 mutated. (**v**) EMSA for *fragment-a* and *fragment-b* with Hth3 and Hth4 mutated. (**w**) EMSA for *fragment-a, fragment-c,* and *fragment-d.* (**x**) EMSA for *fragment-e* with Hth1 mutated. (**y**) EMSA for *fragment-g* with Hth2 mutated.

**Figure S5.**
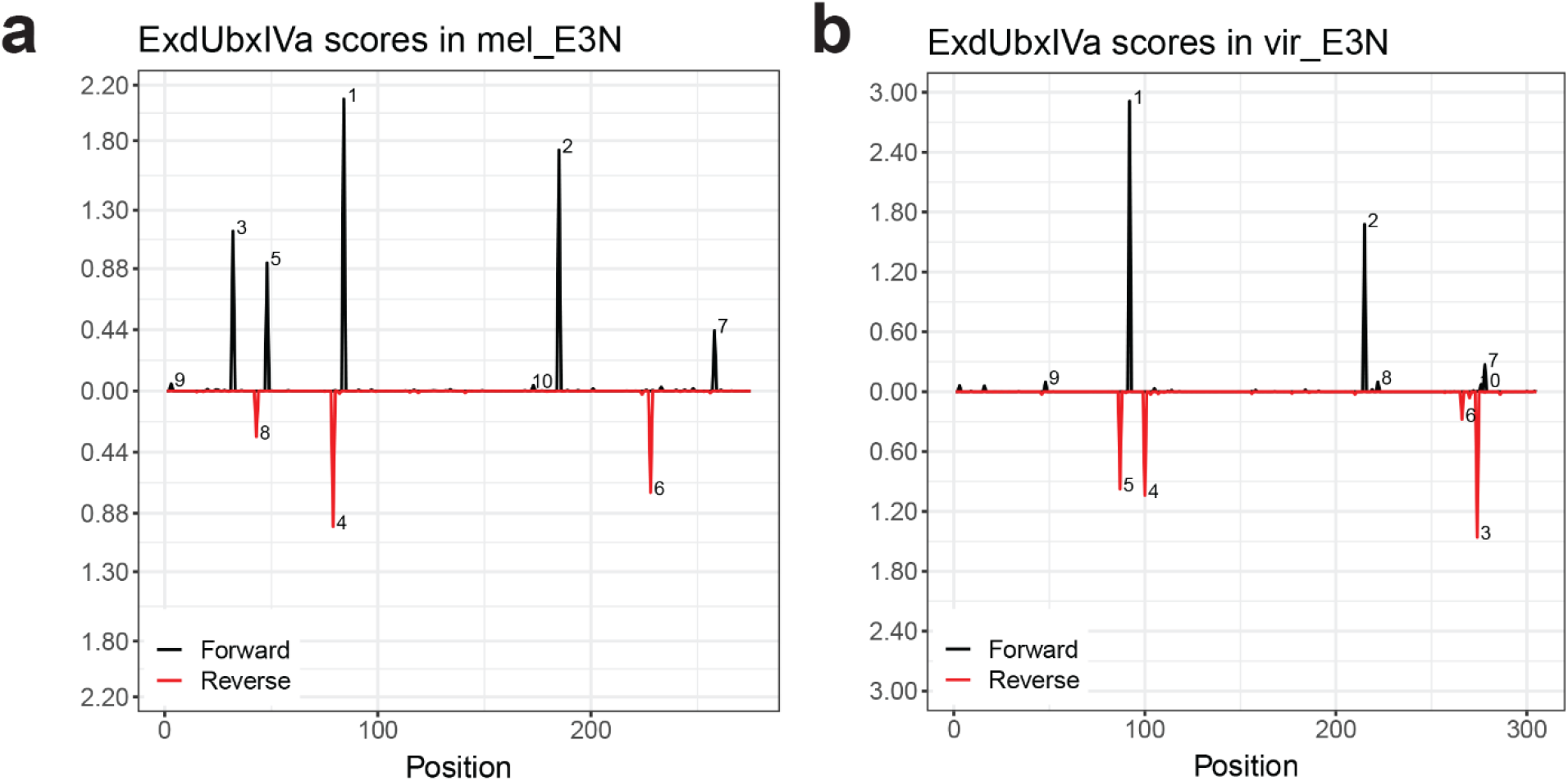
No Read Left Behind (NRLB) total calculated affinities for *D. virilis and D. melanogaster*. (**a and b**) Schematic output from NRLB^40^ shows predicted binding affinity for Exd::Ubx heterodimers across the *E3N* sequence, where black peaks are on the 5’ strand and red peaks respectively on the 3’ strand. Affinity plots are shown for *Drosophila melanogaster* (**a**) and *Drosophila virilis* (**b**).

**Figure S6.**
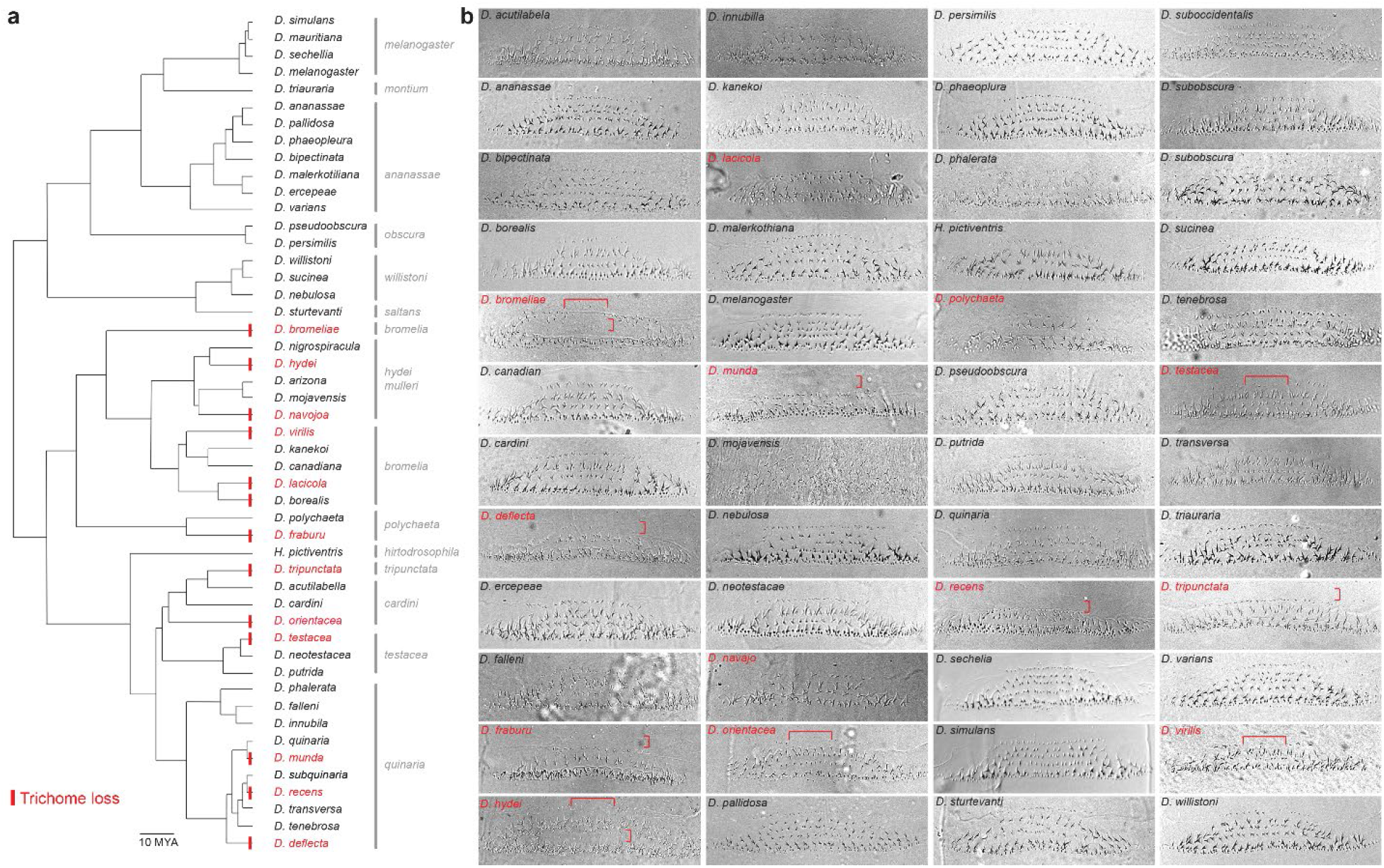
Cuticle preps from 60 *Drosophila* species across approximately 100 million years of evolution. (**a**) *Drosophila e*volutionary tree spanning approximately 150 million years. Red indicates a loss of trichomes. (**b**) Representative cuticle preps for *Drosophila* species. See also Fig. 4.

## Notes

### Competing Interest Statement

The authors have declared no competing interest.

